## Supplemental Results for "Trait salivary alpha-amylase activity levels define the conditions for facilitation by L-DOPA of extinction consolidation"

### Supplementary Results

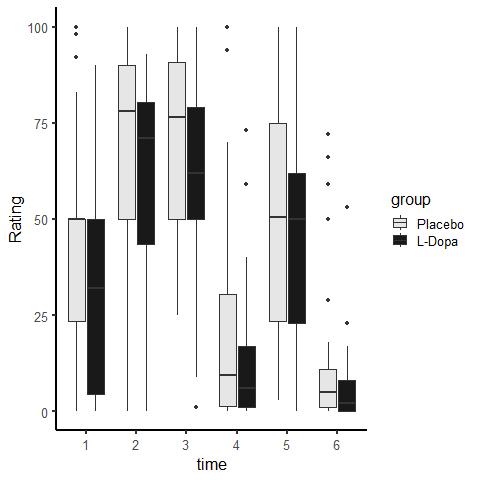

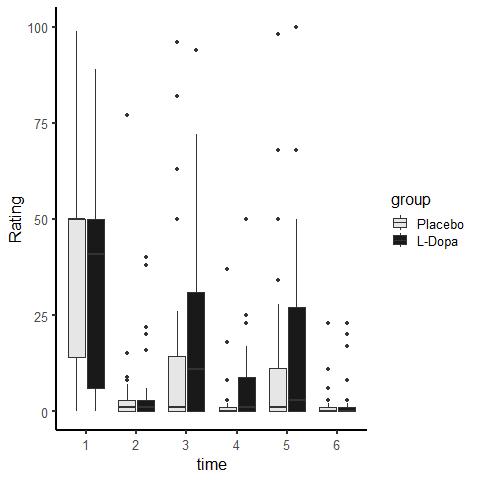

100

75

50

25

0

Fear ratings

Day 1

before after

Day 2

before after

Day 3

before after

Day 1

before after

Day 2

before after

Day 3

before after

CS+

CS-

Placebo

L-DOPA

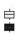

**Supplementary Figure 1. US-expectancy ratings towards CS+ and CS- at all phases.** US-expectancy ratings towards the CS+ (left) and the CS- (right) in both groups throughout all experimental phases. Fear acquisition on day 1 was equally successful in both groups based on subjective ratings, as indicated by a significant effect of stimulus (CS+>CS- ratings after fear acquisition: F_1,68_=246.56, p<.001, generalized η^2^=.65) in the absence of group (placebo/L-DOPA) and stimulus by group effects (ps>.18; n=70). Fear was retrieved at the beginning of extinction on day 2 in both groups (stimulus: F_1,65_=123.58, p<.001, generalized η^2^=.54; group: F_1,65_=0.68, p=.41; stimulus*group: F_1,65_=4.38, p=.04, generalized η^2^=.04, interaction based on higher ratings towards the CS+ in the placebo group: two-sample t-test: t_65_ =2.18, p=.03, CI 95% [-23.91 -1.04], two-sample t-test CS-: p=.26; n=67) and successfully extinguished on day 2 (stimulus: F_1,65_=20.34, p<.001, generalized η^2^=.12; group: F_1,65_=0.55, p=.46; stimulus*group: F_1,65_=4.37, p=.04, generalized η^2^=.03, two-sample t-tests between groups for CS+ and CS- ps>.10; n=67). Administration of L-DOPA on day 2 did not result in a significant group difference on day 3 based on ratings before or after test (group and stimulus*group effects: ps>.06). Data is presented as median ± 1.5*inter quartile range (IQR).

|  | L-DOPA  mean (SD) | Placebo  mean (SD) | t-value / W in Wilcoxon rank test / Anova F-value | p-value |
| --- | --- | --- | --- | --- |
| Age (years) | 30.3 (4.00) (n=35) | 29.2 (4.28) (n=35) | W = 734.5 | .15 |
| BMI (kg/m^2^) | 24.8 (3.55) (n=35) | 24.1 (3.31) (n=35) | W = 600 | .58 |
| STAI-T | 31.3 (6.51) (n=30) | 30.6 (3.89) (n=30) | t = 0.46 | .65 |
| US intensity (mA) | 18.1 (24.3) (n=34) | 14.2 (13.2) (n=34) | W = 524 | .51 |
| US intensity rating (0-10) | 7.26 (1.20) (n=35) | 7.75 (1.07) (n=34) | W = 468 | .12 |
| GHQ | 13.3 (6.43) (n=29) | 12.1 (3.14) (n=30) | W = 457 | .74 |
| AAQ | 24.4 (4.94) (n=35) | 23.6 (3.82) (n=35) | W = 670.5 | 50 |
| STAI-S | D1 begin: 32.3 (6.05) (n=31)  D2 begin: 33.1 (6.54)  D2 end: 29.5 (4.84)  D3 begin: 31.5 (7.92) | D1 begin: 32.6 (6.05) (n=34)  D2 begin: 32.9 (4.05)  D2 end: 30.8 (3.92)  D3 begin: 31.4 (4.70) | Group F = 0.06  Time F = 13.51  Group by Time F = 1.08 | .81  <.001  .36 |
| Rest tiredness (0-10) | D1 rest1: 4.32 (2.23 (n=31)  D1 rest2: 4.61 (1.93)  D2 rest1: 4.90 (2.15)  D2 rest2: 4.52 (2.22)  D2 rest3: 4.16 (2.28)  D2 rest4: 4.26 (2.05)  D3 rest1: 3.58 (2.16) | D1 rest1: 4.70 (2.64) (n=30)  D1 rest2: 4.18 (2.24)  D2 rest1: 4.57 (2.44)  D2 rest2: 3.80 (2.07)  D2 rest3: 3.90 (2.11)  D2 rest4: 3.70 (1.95)  D3 rest1: 3.97 (2.14) | Group F = 0.29  Time F = 2.60  Group by Time F = 1.09 | .59  .02  .36 |

Abbreviations: BMI: body mass index; STAI-T: trait anxiety; US intensity rating on a scale from 0-10 (“I do not feel anything”-“strongest pain imaginable delivered by such an electrode”); GHQ: General Health Questionnaire; AAQ: Acceptance and Action Questionnaire; STAI-S: state anxiety Inventory.

**Supplementary Table 1. Demographic information and questionnaire scores.** There were no significant group differences regarding age, body mass index (BMI), US intensity, subjective rating of the US intensity, or questionnaire scores between L-DOPA and placebo treated groups (all ps>.06). If data was normally distributed (Shapiro-Wilk test) and had equal variance (F-test), a t-test was implemented. Wilcoxon rank test was implemented on data with non-normal distribution. Welch two sample t-test was implemented on data with normal distribution but non-equal variance.

| **ID** | **Group** | **SCR Day 1** | **SCR Day 2** | **SCR Day 3** | **sAA Day 1** | **sAA Day 2** | **sAA Day 3** | **sCORT Day 1** | **sCORT Day 2** | **sCORT Day 3** | **MRI Day 2 task** | **MVP task and rest** |
| --- | --- | --- | --- | --- | --- | --- | --- | --- | --- | --- | --- | --- |
| 1 | Placebo |  |  |  |  |  |  |  |  |  |  |  |
| 2 | L-DOPA |  |  | x |  |  |  |  |  |  |  |  |
| 3 | Placebo |  |  |  |  |  |  |  |  |  |  |  |
| 4 | L-DOPA |  | x |  |  |  |  |  |  |  |  |  |
| 5 | Placebo |  |  |  |  |  |  |  |  |  |  |  |
| 6 | L-DOPA |  |  |  |  |  |  |  |  |  | x | x |
| 7 | L-DOPA |  | x | x |  |  |  |  |  |  |  |  |
| 8 | L-DOPA |  |  |  |  |  |  |  |  |  |  |  |
| 9 | L-DOPA |  | x | x |  |  | x |  |  | x | x | x |
| 10 | L-DOPA | x | x | x |  |  | x |  |  | x | x | x |
| 11 | L-DOPA |  |  | x |  |  |  |  |  |  |  |  |
| 12 | L-DOPA |  |  |  |  |  |  |  |  |  |  |  |
| 13 | L-DOPA |  | x |  |  |  |  |  |  |  |  |  |
| 14 | L-DOPA |  |  |  |  |  |  |  |  |  |  |  |
| 15 | L-DOPA |  | x | x |  |  |  |  |  |  |  |  |
| 16 | L-DOPA |  |  |  |  |  |  |  |  |  |  |  |
| 17 | L-DOPA |  |  |  |  |  |  |  |  |  |  |  |
| 18 | L-DOPA |  |  |  |  |  |  |  |  |  |  |  |
| 19 | Placebo |  |  |  |  |  |  |  |  |  |  |  |
| 20 | L-DOPA |  | x |  |  |  |  |  |  |  |  |  |
| 21 | Placebo |  |  |  |  |  |  |  |  |  |  |  |
| 22 | Placebo |  |  |  |  |  |  |  |  |  |  |  |
| 23 | L-DOPA |  |  |  |  |  |  |  |  |  |  |  |
| 24 | L-DOPA |  | x | x |  |  |  |  |  |  |  |  |
| 25 | Placebo |  |  |  |  |  |  |  |  |  |  |  |
| 26 | Placebo |  |  |  |  |  |  |  |  |  |  |  |
| 27 | L-DOPA |  |  |  | (1/2) |  | (1/2) | (1/2) |  | (1/2) |  |  |
| 28 | L-DOPA |  |  |  | x | (5/7) | (1/2) | x | (2/7) | (1/2) |  |  |
| 29 | L-DOPA |  |  |  |  | (3/7) | (1/2) |  | (2/7) | (1/2) |  |  |
| 30 | Placebo | x |  |  |  |  | (1/2) |  |  |  | x | x |
| 31 | L-DOPA |  | x | x |  | (3/7) |  |  | (3/7) |  | x | x |
| 32 | L-DOPA |  |  |  |  |  |  |  |  |  |  |  |
| 33 | L-DOPA | x |  |  |  |  |  |  |  |  |  |  |
| 34 | Placebo |  |  |  |  |  |  |  |  |  |  |  |
| 35 | Placebo |  |  |  |  |  |  |  |  |  |  |  |
| 36 | Placebo |  |  |  |  | (5/7) |  |  | (2/7) |  |  | x |
| 37 | Placebo |  |  |  | (1/2) |  |  |  |  |  |  |  |
| 38 | L-DOPA |  |  | x |  |  |  |  |  |  |  |  |
| 39 | Placebo |  |  |  |  |  |  |  |  |  |  |  |
| 40 | Placebo |  | x |  |  |  |  |  |  |  |  |  |
| 41 | Placebo |  |  |  |  |  |  |  |  |  |  |  |
| 42 | L-DOPA |  | x |  |  |  |  |  |  |  |  |  |
| 43 | Placebo |  |  |  |  |  |  |  |  |  |  |  |
| 44 | L-DOPA |  |  |  |  |  |  |  |  |  |  |  |
| 45 | Placebo |  |  |  |  |  |  |  |  |  |  |  |
| 46 | Placebo |  | x |  |  |  | (1/2) |  |  | (1/2) |  |  |
| 47 | Placebo |  |  |  |  | (3/7) |  |  |  |  |  |  |
| 48 | Placebo |  |  |  | (1/2) | (5/7) | (2/2) |  | (4/7) | x |  |  |
| 49 | Placebo |  | x | x |  |  |  |  |  |  |  |  |
| 50 | Placebo |  |  |  |  |  |  |  |  |  |  |  |
| 51 | Placebo |  |  |  |  |  |  |  |  |  |  |  |
| 52 | Placebo |  |  |  |  | (4/7) | (1/2) |  | (2/7) |  | x | x |
| 53 | L-DOPA |  | x | x |  |  |  |  |  |  |  |  |
| 54 | L-DOPA |  |  |  |  | (1/7) |  |  |  |  |  |  |
| 55 | Placebo |  | x |  |  |  |  |  |  |  | x | x |
| 56 | L-DOPA |  |  |  | (1/2) | (1/7) |  |  | (1/7) |  | x | x |
| 57 | Placebo |  | x | x |  | x | x |  | x | x | x | x |
| 58 | Placebo | x |  |  | (1/2) |  | (1/2) | (1/2) |  |  |  |  |
| 59 | Placebo |  | x | x |  |  |  |  |  |  |  |  |
| 60 | Placebo |  | x | x |  |  |  |  |  |  |  |  |
| 61 | Placebo |  | x | x |  |  |  |  |  |  |  |  |
| 62 | Placebo |  |  |  |  |  |  |  |  |  |  |  |
| 63 | L-DOPA |  |  |  |  |  |  |  |  |  |  | x |
| 64 | L-DOPA |  |  |  |  |  |  |  |  |  |  |  |
| 65 | L-DOPA |  |  |  | (1/2) | (2/7) | (1/2) |  | (2/7) |  |  |  |
| 66 | Placebo |  |  |  |  |  |  |  |  |  |  | x |
| 67 | Placebo |  |  |  | x | x |  |  | x | x |  |  |
| 68 | Placebo |  |  |  | (1/2) | (3/7) | x | (1/2) | (2/7) | (1/2) | x | x |
| 69 | L-DOPA |  |  |  |  | (3/7) |  |  |  |  |  |  |
| 70 | L-DOPA |  |  |  |  |  |  |  |  |  |  |  |

**Supplementary Table 2. Full exclusion list for all experimental days.** List of all exclusions in SCR data, sAA and sCORT data, and MRI data. x=no data available or excluded, partial data missing in saliva samples presented as (x missing out of/total sample size per day). Number of participants included in separate analyses or analyses comprising several measures are listed at the appropriate paragraphs in the Results part.

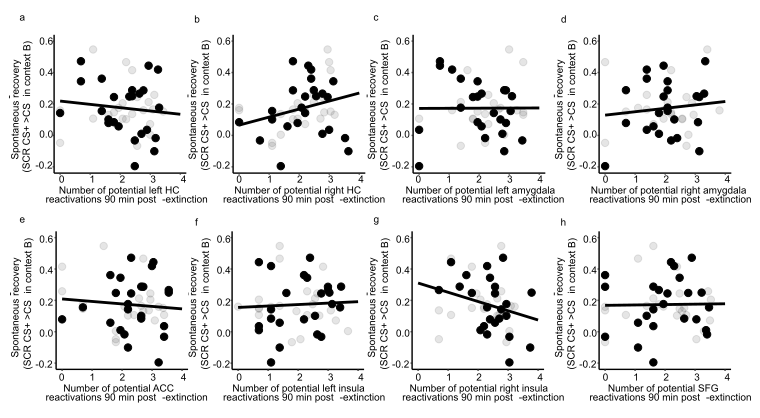

**Supplementary Figure 2. Spontaneous CS+ offset-related pattern reactivations 90 min post extinction on day 2 in control regions do not predict CR (SCR CS+>CS-) at test on day 3.** When including all regions’ number of reactivations, together with the number of reactivations in the vmPFC, into a regression model, the number of CS+ offset-related pattern reactivations in any other region except for the vmPFC did not predict CRs at test (spontaneous recovery in context B) (multiple linear regression: vmPFC: β=-.07, p=.036; all other ps>.11; n=46).

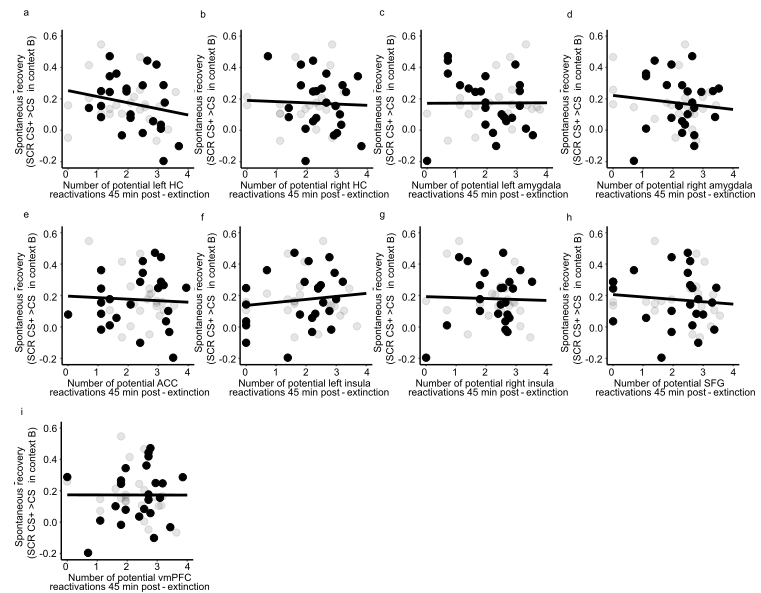
 **Supplementary Figure 3. Spontaneous CS+ offset-related pattern reactivations 45 min post extinction on day 2.** There was no significant relationship between the number of CS+ offset-related pattern reactivations and CRs at test at 45 min post extinction (multiple linear regression, all ps>.09).

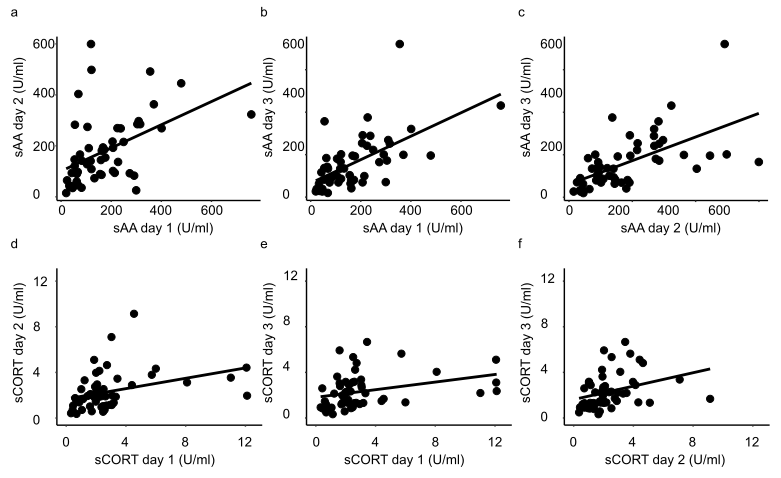

**Supplementary Figure 4**. **Temporal stability of day-to-day sAA and sCORT measures.** We observed highly significant medium- and large-sized correlations in baseline sAA between the three days (day 1-2: R=.46, day 1-3: R=.61, day 2-3: R=.60; all ps<.001). For baseline sCORT, we did not observe temporal stability comparable to baseline sAA (day 1-2: R=.53, day 1-3: R=.32, day 2-3: R=.29; all ps>.11; ICC=.03, p=.43).

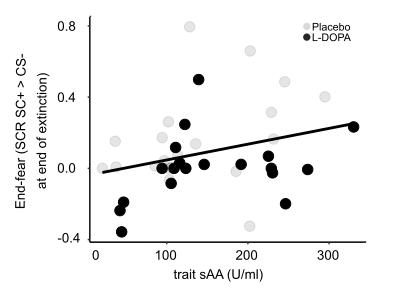

**Supplementary Figure 5. Relationship between trait sAA and end-fear of extinction.** An exploratory analysis revealed that trait sAA levels, operationalized as average of the baseline sAA measures of days 1-3, were associated with high extinction end-fear, that is, poor extinction success (p=.02).

| **Wake up time Day 1** | **Sports on Day 1** | **Smoking on Day 1** |
| --- | --- | --- |
| X-squared = 4.5668, df = 8, p-value = 0.8027 | X-squared = 4.2545, df = 5, p-value = 0.5134 | X-squared = 0, df = 1, p-value = 1 |
| **Wake up time Day 2** | **Sports on Day 2** | **Smoking on Day 2** |
| X-squared = 6.9464, df = 7, p-value = 0.4345 | X-squared = 5.4902, df = 8, p-value = 0.7041 | X-squared = 0, df = 1, p-value = 1 |
| **Wake up time Day 3** | **Sports on Day 3** | **Smoking on Day 3** |
| X-squared = 6.3573, df = 8, p-value = 0.6073 | X-squared = 5.6054, df = 5, p-value = 0.3465 | X-squared = 0.0012039, df = 1, p-value = 0.9723 |

**Supplementary Table 3. Influences of wake up time, sports and smoking on cortisol measurements.** There were no group differences in wake up time (time rounded to full numbers), sports on experimental day (yes/no), and smoking on experimental day (yes/no) on days 1-3 (Pearson's Chi-squared test with Yates' continuity correction). Multiple regression with the first cortisol measurement on each experimental day as the dependent variable and categorical covariates wake up time, sports, smoking, and group revealed an effect of smoking on day 1 (β_smoke_=2.54, SE=1.20, t_62_=2.11, p=.04; n=67, other ps >.23), no effects on day 2 (all ps>.20, n=66), and a significant effect of sports on day 3 on cortisol (β_sports_=-1.25, SE=.42, t_60_=2.99, p=.004; n=60, other ps >.15).

| Side effect | Day 2 L-DOPA | Day 2 Placebo | Day 3 L-DOPA | Day 3 Placebo |
| --- | --- | --- | --- | --- |
| Nausea | 1 | 1 | 1 | 1 |
| Diarrhea | 0 | 0 | 1 | 0 |
| Headache | 5 | 3 | 5 | 4 |
| Dry mouth | 3 | 5 | 2 | 2 |
| Change in sense of taste | 1 | 1 | 0 | 1 |
| Involuntary movement | 0 | 1 | 0 | 0 |
| Tiredness | 14 | 6 | 6 | 2 |
| Sudden sleep attacks | 2 | 1 | 2 | 1 |
| Dizziness | 2 | 2 | 0 | 1 |
| Restlessness | 2 | 2 | 3 | 0 |
| Temporal disorientation | 1 | 1 | 0 | 0 |

**Supplementary Table 4. Side effect questionnaire scores.** Side effects were assessed at the end of day 2 and the beginning of day 3. Eleven out of 18 side effects were reported and are listed here (n out of each group (L-Dopa n=33, placebo n=34) per day). Participants rated the side effect on a scale from “not pronounced”, to “weakly pronounced”, “moderately pronounced”, and “strongly pronounced”. All effects reported here were weakly or moderately pronounced, except for tiredness twice also reported as strongly pronounced. There were no significant group differences in amount of reported side effects (Pearson’s Chi-squared test with simulated p-value (based on 2000 replicates) p=.85).

### Supplementary Methods

**Description of salivary alpha-amylase assays**^1^**:**

Concentration of alpha-amylase in saliva was measured by an enzyme kinetic method: Saliva was processed on a Genesis RSP8/150 liquid handling system (Tecan, Crailsheim, Germany). First, saliva was diluted 1:625 with double-distilled water by the liquid handling system. Twenty microliters of diluted saliva and standard were then transferred into standard transparent 96-well microplates (Roth, Karlsruhe, Germany). Standard was prepared from ‘‘Calibrator f.a.s.’’ solution (Roche Diagnostics, Mannheim, Germany) with concentrations of 326, 163, 81.5, 40.75, 20.38, 10.19, and 5.01 U/l alpha-amylase, respectively, and bidest water as zero standard. After that, 80 ml of substrate reagent (*a*-amylase EPS Sys; Roche Diagnostics, Mannheim, Germany) were pipetted into each well using a multichannel pipette. The microplate containing sample and substrate was then warmed to 37 degrees C by incubation in a waterbath for 90 s. Immediately afterward, a first interference measurement was obtained at a wavelength of 405 nm using a standard ELISA reader (Anthos Labtech HT2, Anthos, Krefeld, Germany). The plate was then incubated for another 5 min at 371C in the waterbath, before a second measurement at 405 nm was taken. Increases in absorbance were calculated for unknowns and standards. Increases of absorbance of diluted samples were transformed to alpha-amylase concentrations using a linear regression calculated for each microplate (Graphpad Prism 4.0c for MacOSX, Graphpad Software, San Diego, CA). The intra and interassay coefficients for amylase were below 9%.

**Description of salivary cortisol assay^1^:**

Saliva samples were frozen and stored at -20 degrees C until analysis. After thawing, salivettes were centrifuged at 3,000 rpm for 5 min, which resulted in a clear supernatant of low viscosity. Salivary concentrations were measured using commercially available chemiluminescence immunoassay with high sensitivity (IBL International, Hamburg, Germany). The intra and interassay coefficients for cortisol were below 9%.

**Description of contingency questionnaire:**

After each experimental day participants filled a contingency questionnaire with following questions:

1) How many symbols did you see?

2) How often do you think the pain stimulus was presented?

3) The presentation of the pain stimulus followed in your opinion a) no systematic, b) no statement possible, c) following systematic: (free text)

4) Did you know when you would receive a pain stimulus? a) yes, b) no, If yes, when?

5) Which room did you see today? (free text)

6) Did you notice anything else you want to report? (free text)

Participants were not excluded based on their responses, as we expect learning even without conscious contingency knowledge. On day 1, 36 out of 70 participants explicitly reported knowledge about the symbol that was associated with the pain stimulus.
